# Metabolic rate is negatively linked to adult survival but does not explain latitudinal differences in songbirds

**DOI:** 10.1101/721779

**Authors:** Andy J. Boyce, James C. Mouton, Penn Lloyd, Blair O. Wolf, Thomas E. Martin

## Abstract

Survival rates vary dramatically among species and predictably across latitudes, but causes of this variation are unclear. The rate of living hypothesis posits that physiological damage from metabolism causes species with faster metabolic rates to exhibit lower survival rates. However, whether increased survival commonly observed in tropical and south temperate latitudes is associated with slower metabolic rate remains unclear. We compared metabolic rates and annual survival rates across 46 species that we measured, and 147 species from literature data across northern, southern, and tropical latitudes. High metabolic rates were associated with lower survival but latitude had substantial direct effects on survival independent of metabolism. The inability of metabolic rate to explain latitudinal variation in survival suggests 1) that species may evolve physiological mechanisms that mitigate physiological damage from cellular metabolism, and 2) a larger role of extrinsic environmental, rather than intrinsic metabolic, causes of latitudinal differences in mortality.

## INTRODUCTION

Adult survival rate varies extensively among species and is a major influence on fitness, demography and life-history evolution (Ashmole 1963; Stearns 1977; Promislow & Harvey 1990; Martin 2015). The rate of living hypothesis has been proposed as a physiological mechanism underlying variation in survival rate. Production of damaging reactive oxygen species (ROS) from metabolism is thought to cause greater oxidative damage and shorter life (Pearl 1928; Harman 1956; Balaban *et al*. 2005; Brys *et al*. 2007; Monaghan *et al*. 2009). Yet, metabolism may be decoupled from senescence because mechanisms to prevent or repair damage (e.g. endogenous antioxidants, mitochondrial membrane composition and telomere dynamics) may coevolve with metabolic rate (Brand 2000; Monaghan & Haussmann 2006; Hulbert *et al*. 2007; Costantini 2008; Salin *et al*. 2015; Skrip & Mcwilliams 2016; Vagasi *et al*. 2018). Furthermore, survival rates may be unrelated to the accumulation of physiological damage entirely. Extrinsic sources of mortality, such as harsh weather or predation, may be more important in shaping variation in survival. Consequently, whether metabolic rate explains variation in adult survival across species remains unclear (Costantini 2008).

Comparative studies show that metabolic rate is negatively correlated with maximum observed lifespan (MLS) in birds and mammals at broad taxonomic scales (Trevelyan *et al*. 1990; Hulbert *et al*. 2007). However, the overriding influence of mass on both lifespan and metabolism obscures the independent effect of metabolic rate on lifespan in such studies (Speakman 2005). Moreover, other comparisons raise questions about this relationship. Bats and birds have higher metabolic rates but are longer-lived than terrestrial mammals of similar size, suggesting that metabolism and lifespan can be decoupled, at least across broad taxonomic groups (Holmes & Austad 1995; Holmes *et al*. 2001; Munshi-South & Wilkinson 2010). Furthermore, measurements of MLS represent exceptional rather than an average across individuals and are sensitive to variation in sample size, recapture probability and quality of record keeping (Krementz *et al*. 1989; Promislow 1993). In addition, MLS is often based on captive individuals that are well-fed and isolated from disease, predation and other extrinsic sources of mortality that are ubiquitous in wild populations (i.e. de Magalhães & Costa 2009). Direct estimates of annual survival rate are not susceptible to these issues.

Adult survival rates vary substantially across latitudes with tropical species generally exhibit higher survival rates than south temperate or especially north temperate species. Yet evidence that metabolic rate underlies variation in adult survival across latitudes is mixed. For example, lower metabolic rates have been found in tropical songbirds (Wikelski *et al*. 2003; Wiersma *et al*. 2007; Londoño *et al*. 2015) and because tropical species are generally longer-lived than temperate relatives (Sandercock *et al*. 2000; Martin 2015; Martin *et al*. 2017), this pattern has been interpreted as evidence that metabolic rate and adult survival are causally linked. However, other studies found no difference in metabolic rates across latitudes in either adult birds (Vleck & Vleck 1979; Bennett & Harvey 1987) or embryos (Martin *et al*. 2013). Furthermore, using latitude as a proxy for survival rate is problematic because survival rates vary extensively among species within latitudes that yield overlap among species between latitudes (reviewed in Martin *et al*. 2017). Metabolic rate and adult survival appeared to be negatively correlated across latitudes in songbirds in one study (Williams *et al*. 2010). However, methods used for estimating survival rates differed between latitudes, which can obscure patterns across sites (Martin *et al*. 2017), and site effects were not included in a statistical test. Within sites, metabolic rate is sometimes negatively linked to adult survival probability (Scholer *et al*. 2019) and sometimes unrelated (Bech *et al*. 2016). Ultimately, studies that directly compare metabolic rates with robust estimates of adult survival from wild populations are needed.

Here, we test whether metabolic rate explains variation in adult survival probability across latitudes. We directly measured resting metabolic rate (RMR) and estimated adult mortality probability for songbirds at North temperate, tropical and South temperate field sites. We also compiled a global database of basal metabolic rate (BMR) and adult survival data for 147 species from the literature. We used phylogenetically-informed path analysis to test whether metabolic rate explained interspecific variation in adult survival within and across latitudes.

## METHODS

### Study species

Passerine birds (songbirds) are a good group in which to examine these issues. Passerines are diverse (∼6,000 species) and show broad ecological and morphological variation (del Hoyo *et al*. 2017). They show large interspecific variation in both metabolic rate (McKechnie & Wolf 2004; Wiersma *et al*. 2007; Londoño *et al*. 2015; McKechnie 2015) and adult survival probability (Johnston *et al*. 1997; Sandercock *et al*. 2000; Martin 2015; Martin *et al*. 2015, 2017).

### Field data

Resting metabolic rate measurements and estimation of adult survival probability were conducted on populations of passerine birds at Kinabalu Park, Sabah, Malaysia (6°N, 116°E), the Coconino National Forest, Arizona, USA (35°N, 111°W) and the Koeberg Nature Reserve, Western Cape, South Africa (34°S, 18°E). Metabolic measurements were performed during the breeding season at both sites (Malaysia; February – June, 2013 – 2016, Arizona; May – July, 2015, South Africa; August – October 2016).

Birds were captured for metabolic measurements by both passive and targeted mist-netting. Breeding females (based on presence of a brood patch) were excluded to minimize disruption of nesting and because the extreme vascularization of the avian brood patch is likely to alter RMR. Birds were transported to the lab and held for 1-2 hrs, depending on mass, to ensure they were post-absorptive during measurements. Birds were watered before and after and returned to point of capture upon completion of metabolic measurements.

Adult survival probability was estimated by banding, resighting and recapturing birds, using the same long-term protocols at Malaysia and Arizona sites (Martin *et al*. 2015). Birds were captured by both passive mist-netting and target-netting for six hours each day beginning at sunrise. Twelve nets were deployed at each netting plot, which were distributed uniformly across accessible areas of each site. Each plot was visited three times at equal intervals over the course of the field season. Birds were marked with unique combinations of one alpha-numeric aluminum band and three color-bands to facilitate individual identification via resighting. In addition to subsequent recaptures, birds were resighted opportunistically each day for the duration of each field season. Similar mark-resight-recapture protocols were used in South Africa (see Lloyd *et al*. 2014). Resulting estimates (see Statistical Analyses) are based on 10 consecutive years of mark-recapture-resighting effort in Borneo, 21 in Arizona, and 7 in South Africa.

### Metabolic measurements

We measured RMR using an open-flow respirometry system similar to that described in Gerson et al. (2015). We used 2 L and 5 L transparent plastic containers (Rubbermaid, Atlanta, GA, USA) as metabolic chambers, depending on the size of the study species. These containers were modified to include incurrent and excurrent air ports, with wire mesh platforms and plastic perches to allow the subject to rest comfortably. The bottom of the chamber contained a 2cm layer of mineral oil to trap moisture and gas associated with feces. Containers were placed inside a large cooler, which was modified to hold an integral peltier device (model AC-162, TE Technology, Traverse City, MI), connected to a temperature controller (Gerson *et al*. 2015) to regulate chamber air temperature. Incurrent air was provided by a high capacity pressure/vacuum pump (model DAA-V515-ED, Gast Manufacturing, Benton Harbor, MI, USA), and was routed through a coil of copper tubing prior to entering the inner chamber to facilitate rapid temperature equilibration. Air flow rates were regulated by mass-flow controllers (Alicat Scientific, Tucson, AZ) with an accuracy of < 2% of the reading and their calibration was checked annually against a factory five-point calibration Alicat mass flow meter used only for this purpose. Flow rates were varied from 2-15 L/min depending on temperature and mass of the study species. Incurrent and excurrent air were both subsampled at rates between 250 and 500 ml/min and CO_2_ and H_2_O were measured using a portable gas-analyzer (LI-COR model LI-840a, Lincoln, NE, USA) zeroed and then spanned against a gas with a known CO_2_ concentration (1854±0.2 ppm). These data were sampled every second and recorded using Expedata (Sable Systems, Las Vegas, NV, USA).

Humidity of incurrent air was regulated using a dew-point generator constructed of three Nalgene bottles connected in series. Air was bubbled through water in the first two bottles, and the third was empty and served as a water trap. The entire device was then submerged in a water bath kept at approximately 10°C by the addition of small ice-packs. This device prevented rapid fluctuations in humidity due to either ambient air temperature or ambient humidity and also prevented condensation occurring in the system. By adjusting water bath temperature and incurrent air pressure, we maintained relative humidity between 50 and 70%, which is within the range of normal conditions at both sites during the breeding season.

Each individual was sampled at multiple temperatures as part of a concurrent study of thermal tolerance. We subsetted data for analysis by selecting the longest continuous period of resting behavior after chamber temperature had reached equilibrium for at least 30 minutes. Subject activity was monitored in real-time via an infrared security camera connected with an external LCD screen. If no period of complete rest greater than two minutes was observed, no data were analyzed for that temperature. We pooled measurements from 27, 30, and 33°C, which are within the thermoneutral zone of most passerines (McKechnie & Wolf 2004; McNab 2009) and selected the lowest measurement for each individual as RMR.

We corrected mass flow rates of humid air, and calculated CO_2_ and H_2_O production using equations in Lighton (2008). Metabolic rate (W) was calculated as in Walsberg and Wolf (1995). CO_2_ production was converted to metabolic energy using a respiratory quotient (RQ) value of 0.71, as suggested for post-absorptive, non-granivorous birds (Gessaman & Nagy 1988).

### Literature data

We compiled basal metabolic rate (BMR) data from the literature, drawing primarily from four manuscripts that use large BMR datasets to investigate allometric and latitudinal variation in avian BMR (McKechnie & Wolf 2004; Wiersma *et al*. 2007; Londoño *et al*. 2015; Bech *et al*. 2016). Estimates of annual adult survival probability were compiled by searching the literature, and were greatly aided by manuscripts containing large literature (Martin 1995; Martin & Clobert 1996) and field (Scholer *et al*. 2019) datasets. Where multiple estimates of either BMR or adult survival probability for a single species were present in the literature, we chose the estimate based on the most recent study.

### Statistical Analyses

For our Malaysia and Arizona field data, we employed Cormack-Jolly-Seber models to estimate apparent annual adult survival (ϕ) and detection probability (*p*) for each species based on live encounters in an open population using program MARK (White & Burnham 1999; Burnham & Anderson 2002). A suite of models were built for each species, allowing parameters to vary based on sex and/or age-structure (time since marking; Pradel *et al*. 1997). Top models were selected based on Akaike’s information criterion (AICc) adjusted for small sample size. Estimates used here are an updated subset of those presented in Martin *et al*. (Martin *et al*. 2015, 2017), where additional methodological details are provided. For our South Africa data, we similarly fit Cormack-Jolly-Seber models but survival was held constant and year was treated as a random effect. Estimates used here are from Lloyd *et al*. (2014).

We used simple linear regression to examine the expected allometric relationship between species mean body mass and metabolic rate for both field RMR and literature BMR datasets. We log-transformed RMR (W), BMR (W), and body mass (g).

We used phylogenetic path analysis (PPA) to examine the causal relationships between mass, metabolic rate, apparent adult survival, and latitude (Hardenberg & Gonzalez-Voyer 2013). PPA uses the d-separation method to test the plausibility that a causal model created the observed data and to compare the relative support of mulitple models. We developed six possible models (Fig. 1) that varied in the depictions of how latitude and metabolic rate influence survival and how latitude influences metabolic rate. We tested the conditional independencies of each model using phylogenetic least-squares regression (PGLS), implemented in the package ‘ape’ (Paradis *et al*. 2004; Popescu *et al*. 2012). We then tested the plausibility of each causal model using Fisher’s C statistic. We used the C statistic Information Criterion with a correction for small sample sizes (CICc) to rank and compute the probability of each causal model given the data and the candidate model set (CIC weight). We then used model averaging to estimate standardized path coefficients for all plausible models (p > 0.05; Anderson *et al*. 2000; Hardenberg & Gonzalez-Voyer 2013). We log-transformed RMR (W), BMR (W), and body mass (g). We followed identical procedures for analysis of both our field RMR data and literature BMR data.

**Figure 1.**
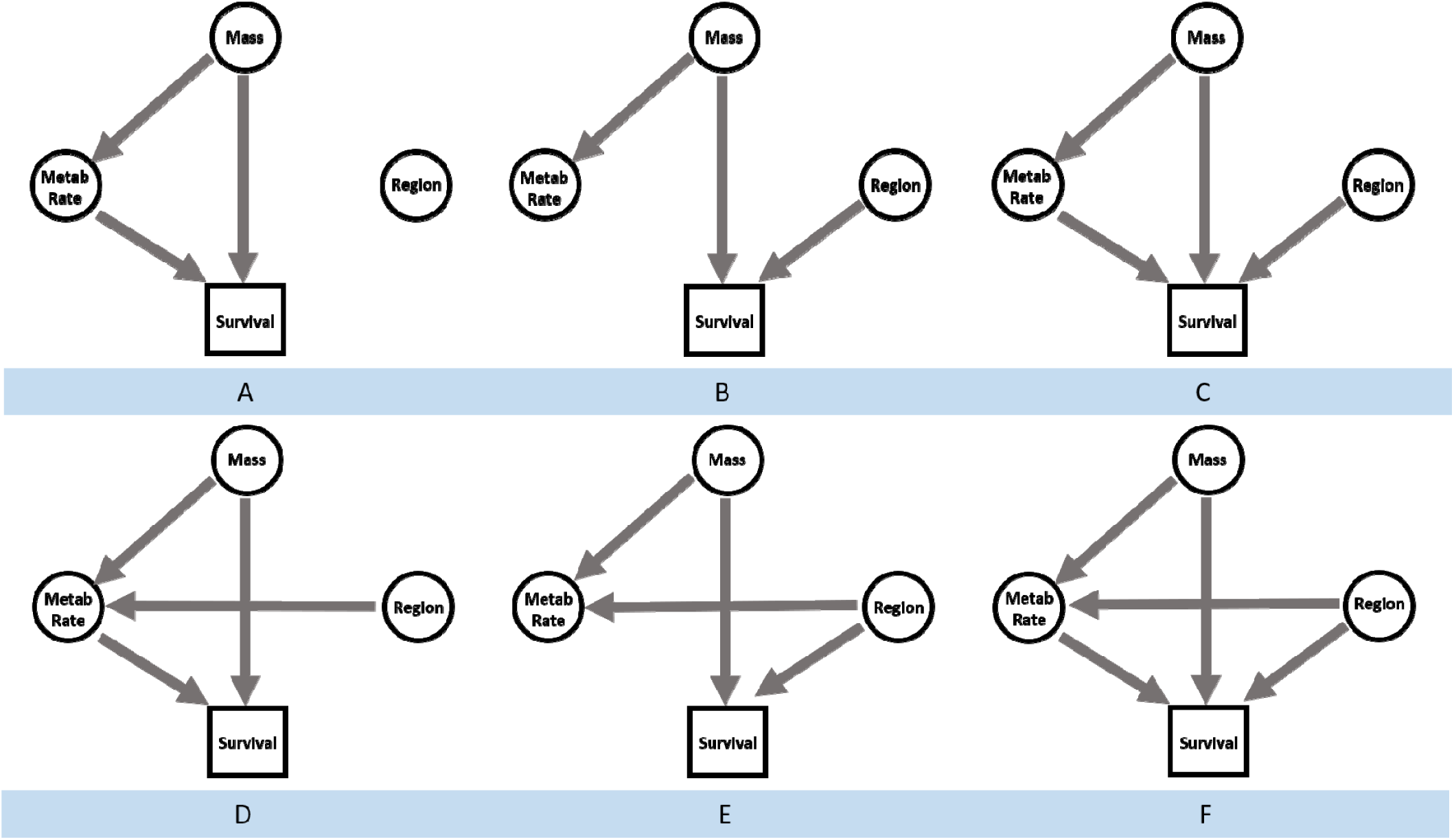
Our candidate model set (A-F) for phylogenetic path analysis. Models visually describe possible causal relationships between body mass, metabolic rate, survival probability and site/region. Metabolic rate is resting metabolic rate (RMR) for field studies and basal metabolic rate (BMR) for literature data.

Phylogenetic trees were acquired from birdtree.org (Jetz *et al*. 2012) using the Hackett backbone (Hackett *et al*. 2008). We sampled a distribution of 1000 trees for each analysis and produced majority-rules consensus trees using Mesquite (Maddison & Maddison 2011). In one case (*Troglodytes aedon*), we include both a tropical (*T. a. musculus*) and temperate (*T. a. aedon*) subspecies in our literature BMR dataset. To facilitate phylogenetic analysis in which duplicate estimates for a single species are problematic, we assigned one estimate to a closely related congener (*T. cobbi*) for tree construction.

Finally, we combined field and literature datasets and tested for differences in allometric relationships between RMR and BMR using a simple linear model. All analyses were performed in R (R Core Team 2015).

## RESULTS

We obtained field-based estimates of adult mortality probability and resting metabolic rate for 47 species; 14 in Arizona, 16 in Malaysia, and 17 in South Africa. We measured RMR in a total of 237 individuals with an average of 5.0 individuals per species (Table 1). Body mass explained the majority of variation in metabolic rates (R^2^ = 0.94, *P* < 0.01, Fig. 2A). The allometric scaling exponent was 0.64, which is consistent with known values for birds (Bennett & Harvey 1987; McKechnie & Wolf 2004). The only plausible model that explained relationships was the full model (Fig. 1F) which described direct causal relationships between RMR and survival, latitude and survival, and latitude and RMR (Fig. 2C, 3A). Adult survival declined with RMR, but was higher in tropical and south temperate species compared with north temperate relatives after accounting for RMR (Fig. 2C, 3A). RMR was slightly lower in tropical species, but higher in the south temperate when compared to north temperate species (Fig. 2C, 3A).

**Table 1.**
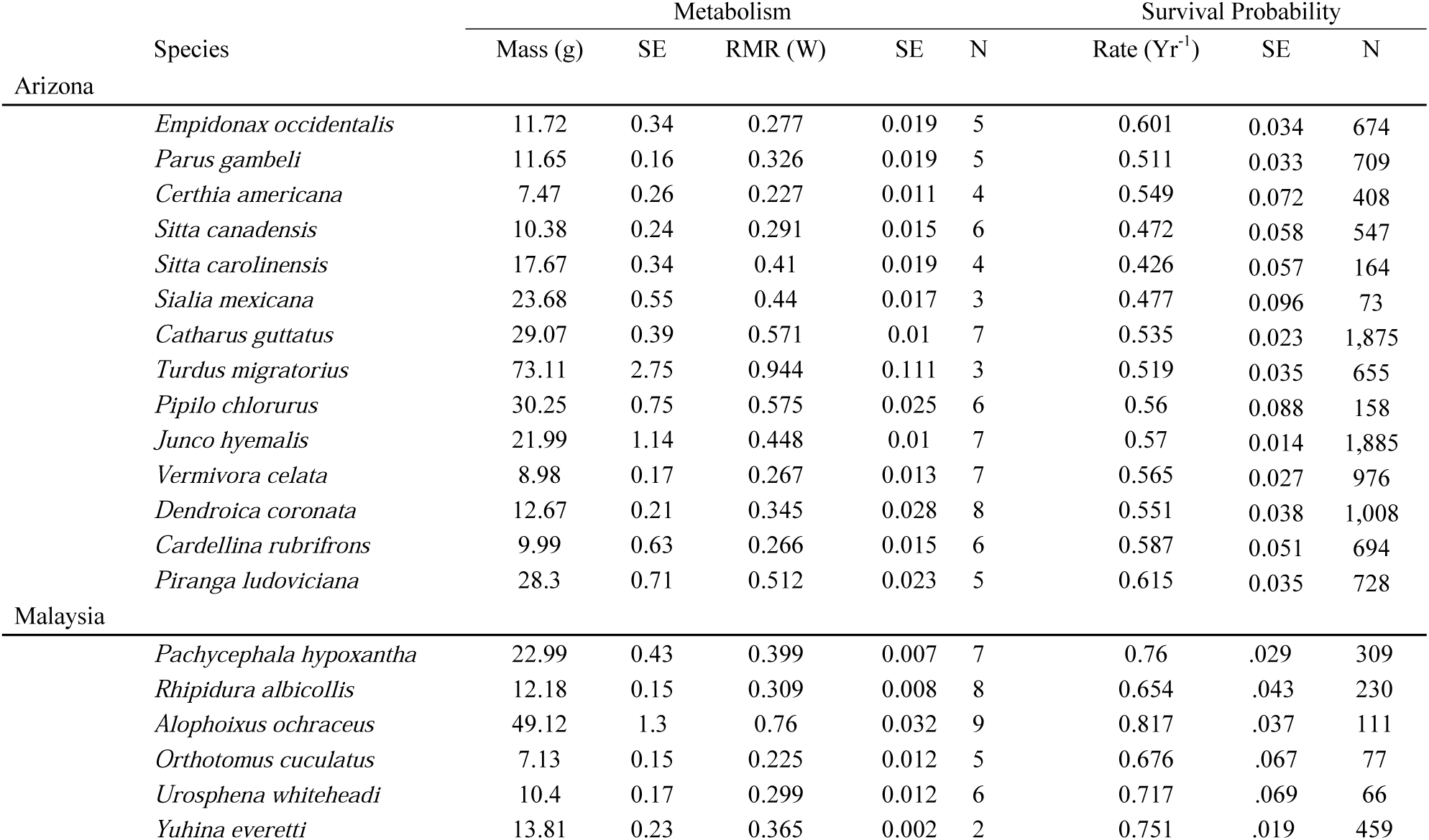
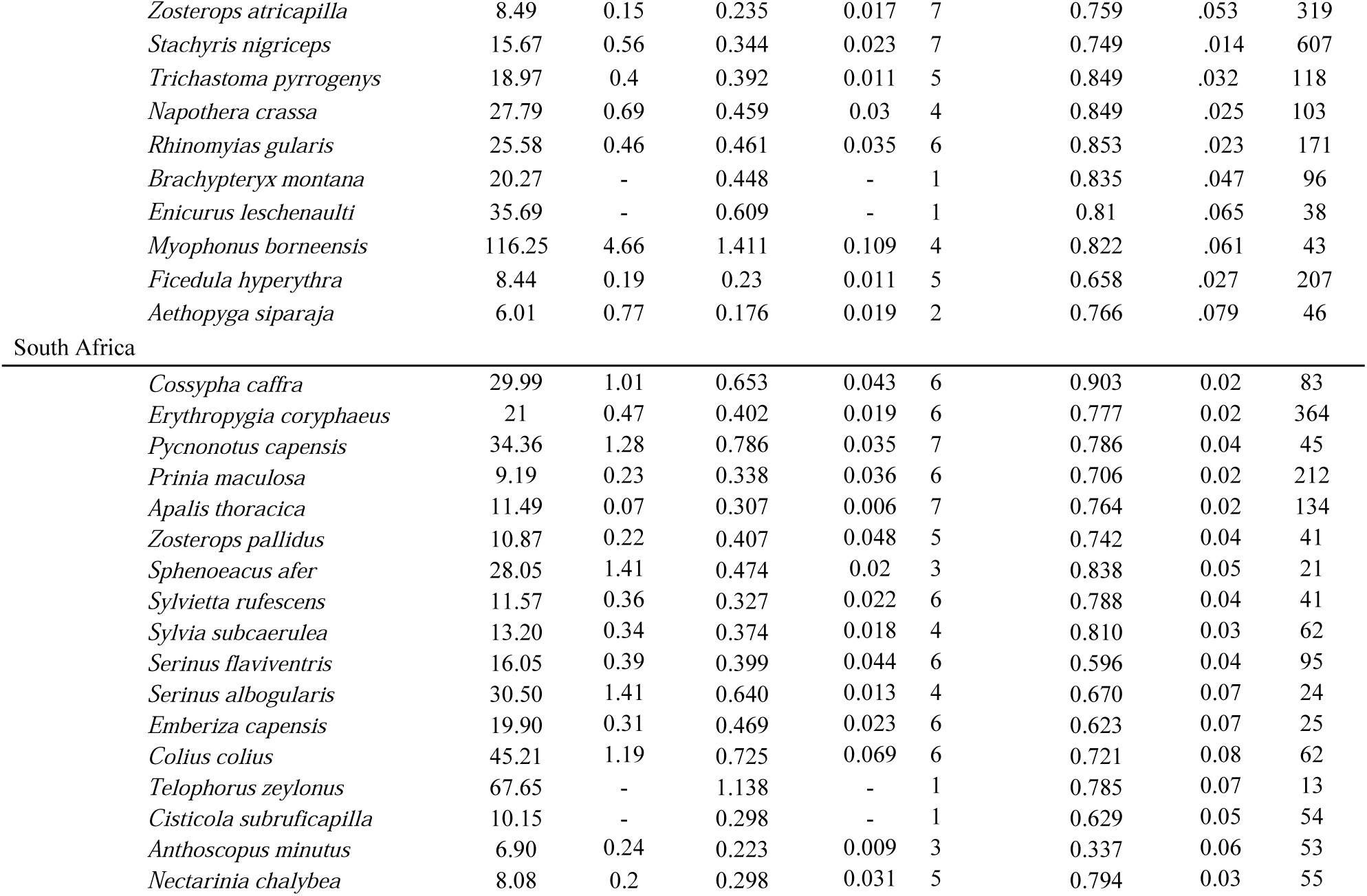
Mean (± SE) values of mass, resting metabolic rate (RMR) and annual mortality probability for species in our field studies. N (sample size) values represent the number of unique individuals measured for metabolic rate and the number of unique individuals marked for annual mortality probability.

| Species | Metabolism |  |  |  |  | Survival Probability |  |  |
| --- | --- | --- | --- | --- | --- | --- | --- | --- |
|  | Mass (g) | SE | RMR (W) | SE | N | Rate (Yr<sup>-1</sup>) | SE | N |
| Arizona |  |  |  |  |  |  |  |  |
| *Empidonax occidentalis* | 11.72 | 0.34 | 0.277 | 0.019 | 5 | 0.601 | 0.034 | 674 |
| *Parus gambeli* | 11.65 | 0.16 | 0.326 | 0.019 | 5 | 0.511 | 0.033 | 709 |
| *Certhia americana* | 7.47 | 0.26 | 0.227 | 0.011 | 4 | 0.549 | 0.072 | 408 |
| *Sitta canadensis* | 10.38 | 0.24 | 0.291 | 0.015 | 6 | 0.472 | 0.058 | 547 |
| *Sitta carolinensis* | 17.67 | 0.34 | 0.41 | 0.019 | 4 | 0.426 | 0.057 | 164 |
| *Sialia mexicana* | 23.68 | 0.55 | 0.44 | 0.017 | 3 | 0.477 | 0.096 | 73 |
| *Catharus guttatus* | 29.07 | 0.39 | 0.571 | 0.01 | 7 | 0.535 | 0.023 | 1,875 |
| *Turdus migratorius* | 73.11 | 2.75 | 0.944 | 0.111 | 3 | 0.519 | 0.035 | 655 |
| *Pipilo chlorurus* | 30.25 | 0.75 | 0.575 | 0.025 | 6 | 0.56 | 0.088 | 158 |
| *Junco hyemalis* | 21.99 | 1.14 | 0.448 | 0.01 | 7 | 0.57 | 0.014 | 1,885 |
| *Vermivora celata* | 8.98 | 0.17 | 0.267 | 0.013 | 7 | 0.565 | 0.027 | 976 |
| *Dendroica coronata* | 12.67 | 0.21 | 0.345 | 0.028 | 8 | 0.551 | 0.038 | 1,008 |
| *Cardellina rubrifrons* | 9.99 | 0.63 | 0.266 | 0.015 | 6 | 0.587 | 0.051 | 694 |
| *Piranga ludoviciana* | 28.3 | 0.71 | 0.512 | 0.023 | 5 | 0.615 | 0.035 | 728 |
| Malaysia |  |  |  |  |  |  |  |  |
| *Pachycephala hypoxantha* | 22.99 | 0.43 | 0.399 | 0.007 | 7 | 0.76 | .029 | 309 |
| *Rhipidura albicollis* | 12.18 | 0.15 | 0.309 | 0.008 | 8 | 0.654 | .043 | 230 |
| *Alophoixus ochraceus* | 49.12 | 1.3 | 0.76 | 0.032 | 9 | 0.817 | .037 | 111 |
| *Orthotomus cuculatus* | 7.13 | 0.15 | 0.225 | 0.012 | 5 | 0.676 | .067 | 77 |
| *Urosphena whiteheadi* | 10.4 | 0.17 | 0.299 | 0.012 | 6 | 0.717 | .069 | 66 |
| *Yuhina everetti* | 13.81 | 0.23 | 0.365 | 0.002 | 2 | 0.751 | .019 | 459 |
| *Zosterops atricapilla* | 8.49 | 0.15 | 0.235 | 0.017 | 7 | 0.759 | .053 | 319 |
| *Stachyris nigriceps* | 15.67 | 0.56 | 0.344 | 0.023 | 7 | 0.749 | .014 | 607 |
| *Trichastoma pyrogenys* | 18.97 | 0.4 | 0.392 | 0.011 | 5 | 0.849 | .032 | 118 |
| *Napothera crassa* | 27.79 | 0.69 | 0.459 | 0.03 | 4 | 0.849 | .025 | 103 |
| *Rhinomyias gularis* | 25.58 | 0.46 | 0.461 | 0.035 | 6 | 0.853 | .023 | 171 |
| *Brachypteryx montana* | 20.27 | - | 0.448 | - | 1 | 0.835 | .047 | 96 |
| *Enicurus leschenaulti* | 35.69 | - | 0.609 | - | 1 | 0.81 | .065 | 38 |
| *Myophonus borneensis* | 116.25 | 4.66 | 1.411 | 0.109 | 4 | 0.822 | .061 | 43 |
| *Ficedula hyperythra* | 8.44 | 0.19 | 0.23 | 0.011 | 5 | 0.658 | .027 | 207 |
| *Aethopyga siparaja* | 6.01 | 0.77 | 0.176 | 0.019 | 2 | 0.766 | .079 | 46 |
| South Africa |  |  |  |  |  |  |  |  |
| *Cossypha caffra* | 29.99 | 1.01 | 0.653 | 0.043 | 6 | 0.903 | 0.02 | 83 |
| *Erythropgia coryphaeus* | 21 | 0.47 | 0.402 | 0.019 | 6 | 0.777 | 0.02 | 364 |
| *Pycnonotus capensis* | 34.36 | 1.28 | 0.786 | 0.035 | 7 | 0.786 | 0.04 | 45 |
| *Prinia maculosa* | 9.19 | 0.23 | 0.338 | 0.036 | 6 | 0.706 | 0.02 | 212 |
| *Apalis thoracica* | 11.49 | 0.07 | 0.307 | 0.006 | 7 | 0.764 | 0.02 | 134 |
| *Zosterops pallidus* | 10.87 | 0.22 | 0.407 | 0.048 | 5 | 0.742 | 0.04 | 41 |
| *Sphenoeacus afer* | 28.05 | 1.41 | 0.474 | 0.02 | 3 | 0.838 | 0.05 | 21 |
| *Sylvietta rufescens* | 11.57 | 0.36 | 0.327 | 0.022 | 6 | 0.788 | 0.04 | 41 |
| *Sylvia subcaerulea* | 13.20 | 0.34 | 0.374 | 0.018 | 4 | 0.810 | 0.03 | 62 |
| *Serinus flaviventris* | 16.05 | 0.39 | 0.399 | 0.044 | 6 | 0.596 | 0.04 | 95 |
| *Serinus albogularis* | 30.50 | 1.41 | 0.640 | 0.013 | 4 | 0.670 | 0.07 | 24 |
| *Emberiza capensis* | 19.90 | 0.31 | 0.469 | 0.023 | 6 | 0.623 | 0.07 | 25 |
| *Colius colius* | 45.21 | 1.19 | 0.725 | 0.069 | 6 | 0.721 | 0.08 | 62 |
| *Telophorus zeylonus* | 67.65 | - | 1.138 | - | 1 | 0.785 | 0.07 | 13 |
| *Cisticola subruficapilla* | 10.15 | - | 0.298 | - | 1 | 0.629 | 0.05 | 54 |
| *Anthoscopus minutus* | 6.90 | 0.24 | 0.223 | 0.009 | 3 | 0.337 | 0.06 | 53 |
| *Nectarinia chalybea* | 8.08 | 0.2 | 0.298 | 0.031 | 5 | 0.794 | 0.03 | 55 |

**Figure 2.**
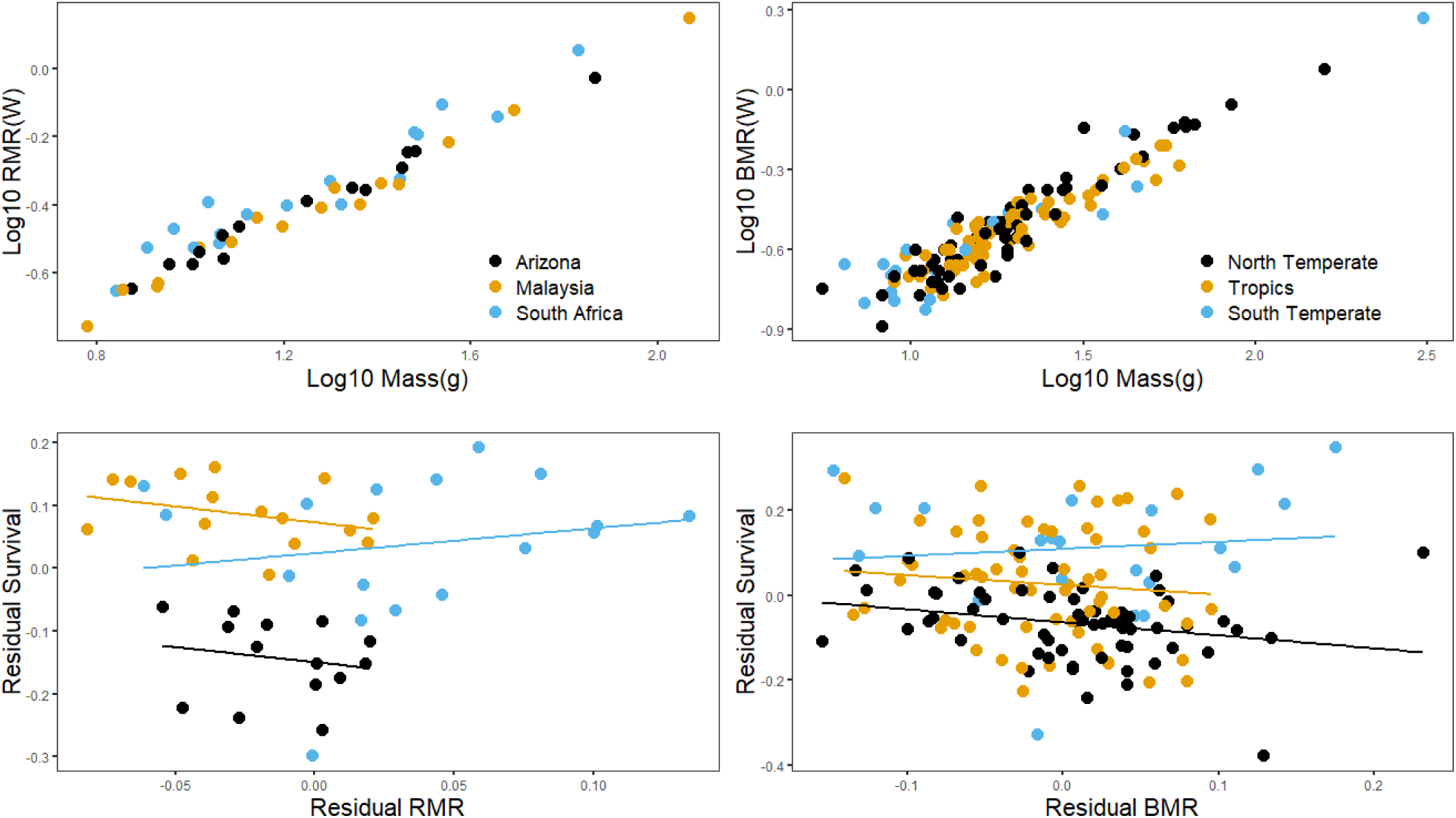
Allometric scaling of resting metabolic rate (A) and basal metabolic rate (B) and body mass for our field and literature datasets. The relationship between resting (C) and basal (D) metabolic rate and adult survival for species in our field and literature datasets. Both variables are residual values controlled for body mass. Each point represents mean values for an individual species. Throughout the plots, north temperate species are in black, tropical species in yellow and south temperate species in blue.

**Figure 3.**
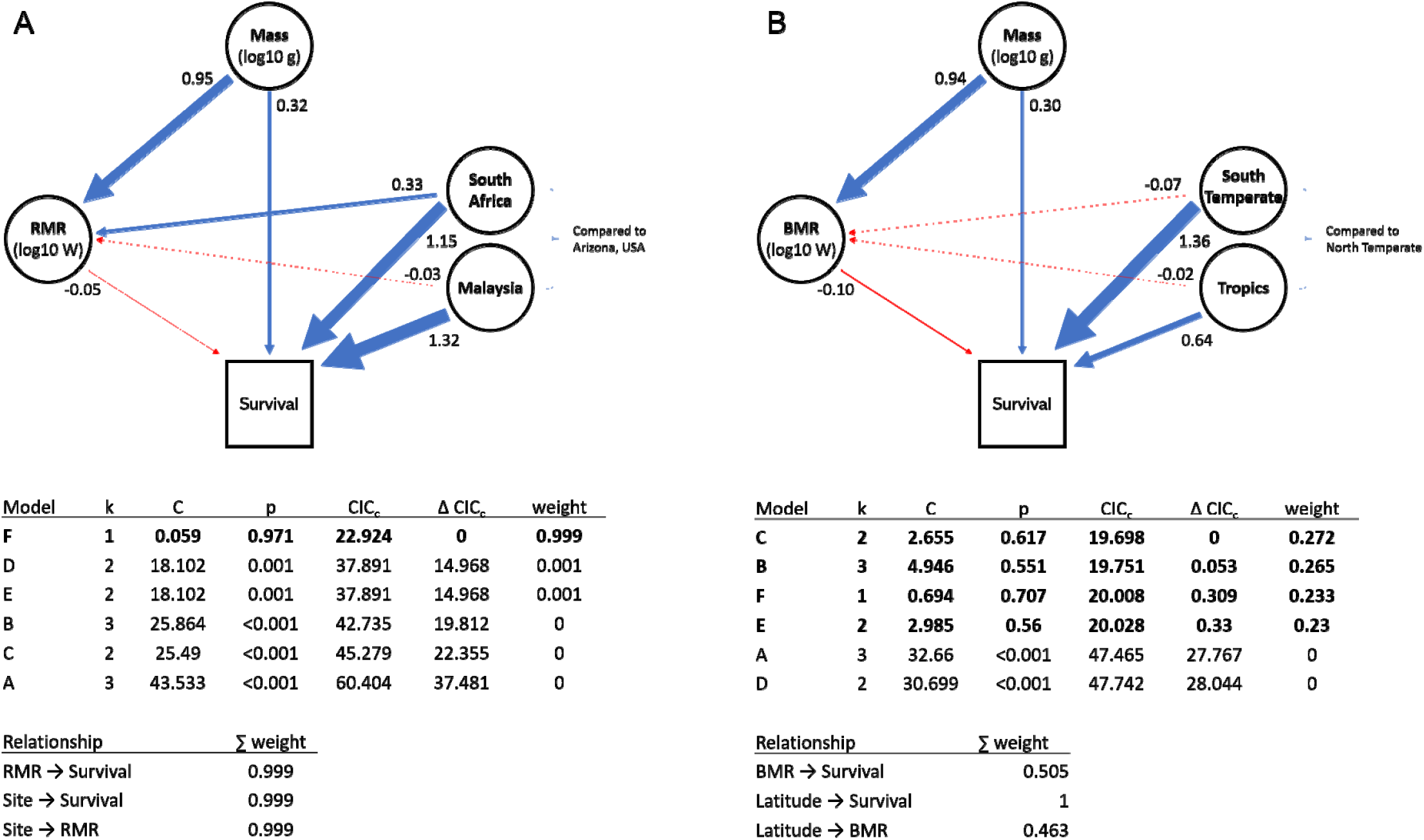
Visual and tabular representation of the causal relationships between body mass, latitude, metabolic rate and adult survival probability for our field and literature datasets. Blue lines represent positive relationships and negative relationships are in red. Arrows point in the direction of causality (from cause to effect). The width of each arrow is proportional to the effect size in number of standard deviations in variation explained according to weighted model averaging. Arrows are only present for relationships present in a subset of plausible models. Bolded models in CIC_c_ tables are those which are plausible given the data (α = 0.05).

Our compiled literature data for BMR and adult survival probability included 62 temperate, 64 tropical, and 21 south temperate species (Supplementary Table 1). Similar to our field data, body mass explained extensive variation in metabolic rates (R^2^ = 0.86, *P* < 0.01, Fig 2B). The allometric scaling exponent was 0.66, nearly identical to our RMR estimate above. Four models could have plausibly created the observed BMR and survival data from the literature (Fig. 1, 3B). Among these, a direct effect of latitude on adult survival was strongly supported (CICc weight = 1.0). A negative relationship between BMR and survival also had some support (CICc weight = 0.505), as well as latitudinal differences in BMR (CICc weight = 0.463). Adult survival declined with BMR and was higher in tropical and south temperate species than north temperate relatives after accounting for BMR (Fig. 2D, 3B). BMR was slightly lower in tropical and south temperate species than in north temperate species (Fig. 2D, 3B).

RMR was 14.7% higher than BMR (*P* < 0.01), but the allometric relationship between mass and metabolism did not differ for (RMR) and basal (BMR) metabolic rates (*P* = 0.51, Fig. 7).

## DISCUSSION

Broad tests of metabolism and annual survival probability across diverse species have been lacking, despite a long history of their possible association ((Pearl 1928; Harman 1956; Balaban *et al*. 2005; Brys *et al*. 2007; Monaghan *et al*. 2009) that has been challenged by cross-taxa comparisons (i.e., Holmes & Austad 1995; Holmes *et al*. 2001; Munshi-South & Wilkinson 2010). Our field data on RMR and those from the literature on BMR included 193 species across the world and yielded results that were largely in agreement between the two datasets. In particular, the results suggest that metabolic rate is associated with adult survival, but most of the variation in adult survival probability among latitudes is independent of metabolic rate (Fig. 2, 4). Our results provide some support for a possible role of the rate-of-living hypothesis within latitudes. However, more significantly, our results parallel those of comparisons across taxonomic groups (i.e., Holmes & Austad 1995; Holmes *et al*. 2001; Munshi-South & Wilkinson 2010) in suggesting that metabolic rate is not the primary driver of broader global patterns of survival probability.

**Figure 4.**
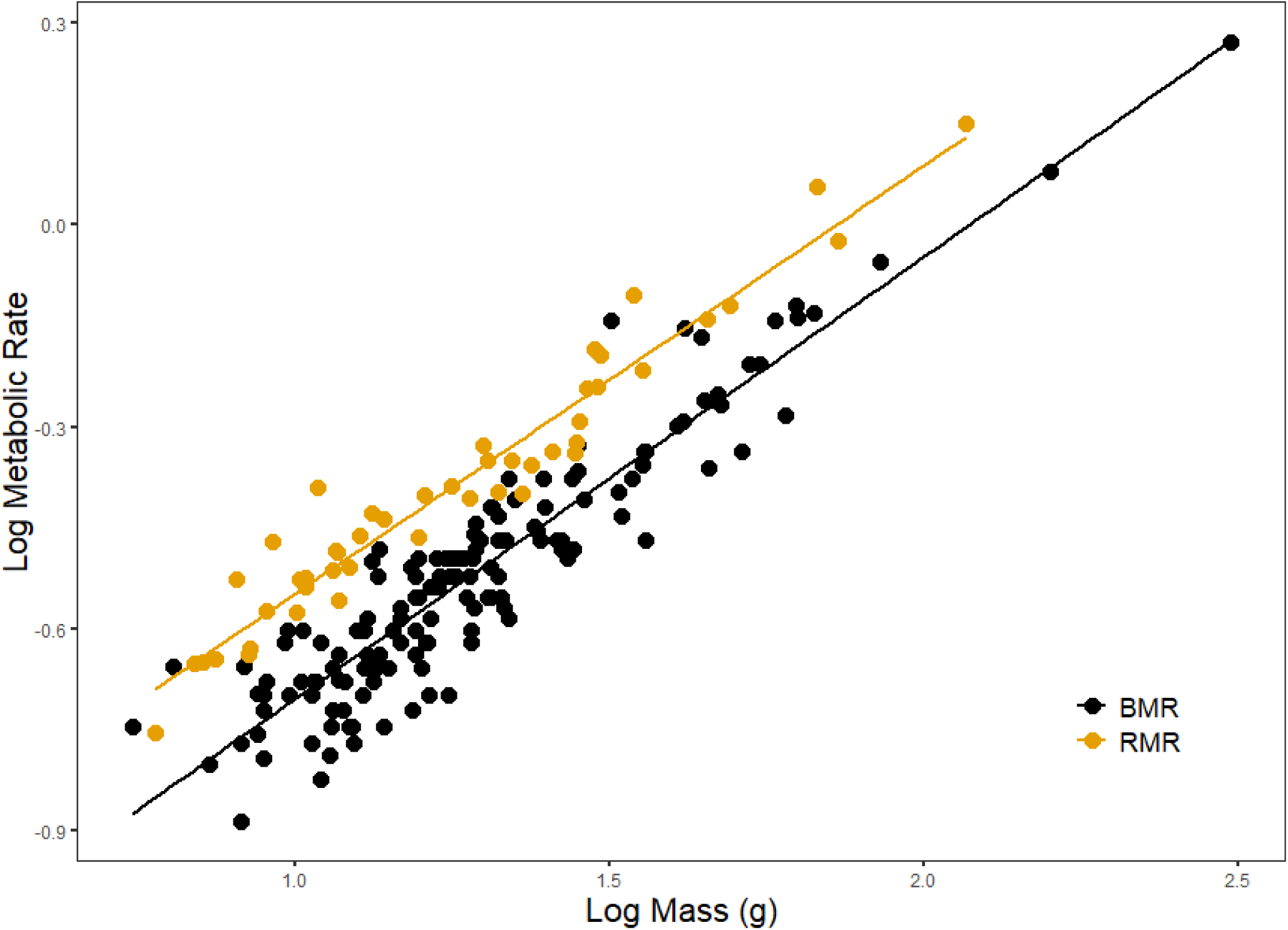
Allometric scaling relationships of RMR and BMR. Each point represents individual species-mean values for body mass and metabolic rate. RMR measurements are in yellow, BMR points are in black.

We found that while metabolic rates were reduced in tropical compared with north temperate species, results were mixed when comparing south temperate to north temperate species (Fig. 2A-B, 3). Lower metabolism in tropical species fits with results of previous studies (Wiersma *et al*. 2007; Londoño *et al*. 2015). However, the difference in metabolism across latitudes is only a small, statistical difference (as also found by Wiersma *et al*. 2007, Fig 1; Londono *et al*. 2015), whereas adult survival probability shows a consistently large difference between the north temperate and tropical latitudes (Martin *et al*. 2017). The small difference in metabolism and larger difference in survival fits with our results that much of the difference in survival probabilities between latitudes is not related to metabolism.

Consideration of metabolism and survival in south temperate regions has been limited (but see Bech *et al*. 2016). However, inclusion of this region provides critical additional insight. Our field dataset indicated higher resting metabolic rate in South African species compared with Arizona, but support for the significance of this relationship was relatively weak (Fig. 3A). Nonetheless, the resting metabolic rates do not fit the pace-of-life hypothesis given that the Southern African species have substantially higher survival rates than Arizona species (Peach *et al*. 2001; Lloyd *et al*. 2014). In contrast, BMR was reduced in south temperate compared with north temperate species based on literature data (Fig. 3B), and this relationship was strongly supported. The high survival rates of south temperate species (Lloyd *et al*. 2014) taken together with their lower BMR rates then fits with the pace-of-life hypothesis (Wiersma *et al*. 2007; Healy *et al*. 2019). This inconsistency between RMR and BMR together with earlier work in both adult birds (Vleck & Vleck 1979; Bennett & Harvey 1987) and embryos (Martin *et al*. 2013) that show no latitudinal difference in metabolism, suggests that metabolic rate is a minor influence on latitudinal variation in survival rates relative to other factors associated with latitude.

Latitudinal variation in avian mortality rates may be driven primarily by differences in extrinsic mortality probability. Extrinsic mortality is thought to account for 80-95% of all mortality for birds with total annual mortality rates similar to those in our study (Ricklefs 1998). Thus, variation in extrinsic mortality is likely to have a much larger effect on total mortality rates than intrinsic physiological differences. However, reduced metabolic costs to survival should be favored in populations with low extrinsic adult mortality, meaning extrinsic and intrinsic mortality rates should be correlated (reviewed in (Charlesworth 1994, 2000). Indeed, actuarial (Promislow 1991; Ricklefs 1998, 2000) and experimental (Stearns *et al*. 2000) studies across taxa suggest intrinsic mortality rate increases with extrinsic mortality rate (e.g. weather, predation). However, the proportion of deaths from intrinsic sources are greater when overall mortality rates are low, suggesting that adaptations to slow the rate of aging are limited, such that extrinsic and intrinsic rates may become increasingly decoupled as extrinsic mortality declines (Ricklefs & Scheuerlein 2001). Furthermore, the onset of senescence is commonly delayed until well after the age of maturity (Promislow 1991), suggesting that intrinsic and extrinsic mortality rates may also be unrelated when extrinsic mortality is very high. Ultimately, high adult mortality rates in temperate birds may reflect high rates of extrinsic mortality imposed by abiotic factors (MacArthur 1972) that better explain latitudinal differences in survival (Martin 2002, 2015; Martin *et al*. 2015).

The absence of a strong relationship between metabolic rate and adult survival across latitudes does not discount the possibility that physiological damage from cellular metabolism contributes to adult survival rates and life-history tradeoffs. On the contrary, a causal relationship between metabolism and survival was supported (Fig. 3) and the predicted negative associations were observed within most regions using both RMR and BMR datasets. Moreover, increased investment in mechanisms to mitigate damage, such as endogenous production of antioxidants (Parolini *et al*. 2017) or mitochondrial membrane composition (Hulbert *et al*. 2007), can reduce damage from cellular metabolism. If tropical species invest in these mechanisms with allocation costs for growth or reproduction, such a tradeoff could explain the longer life and slower life-history strategies of tropical species despite broadly similar metabolic rates across latitudes. Yet, ultimately, investment in such mechanisms only makes sense if extrinsic mortality is low.

BMR and RMR are the most easily measured and comparable metrics of energy expenditure in wild organisms. However, these measures only encompass minimal energy expenditure to sustain life and thus exclude energy allocated to essential activities such as reproduction, thermoregulation, locomotion and digestion. Physiological damage from metabolism may be more tightly linked to measures of total energy expenditure that describe all energetic expenditures in free-living organisms. Measurements of total energy expenditure, such as field metabolic rate (FMR) or daily energy expenditure (DEE) are comparatively rare in the literature, especially for tropical species (McKechnie 2015), but do show a relationship with adult survival probability in the temperate zone (Martin 2014). BMR and RMR are strongly correlated with each other (Fig. 4), and with measures of total energy expenditure across species (Daan *et al*. 1990; Auer *et al*. 2017), making BMR and RMR reasonable but imperfect proxies for total energy expenditure. Future studies should examine the relationship between FMR and adult survival within and across latitudes.

Our study provides support for some role for the rate-of-living hypothesis within latitudes while also suggesting that it is only explains a small amount of the variation in survival within latitudes and is unable to explain differences between latitudes. This contradiction provides obvious opportunity for future studies. South temperate and tropical birds have longer developmental periods and parents invest more energy per-offspring compared with temperate species (Martin 1996, 2015; Martin *et al*. 2011; Gill & Haggerty 2012). These differences may facilitate longer life in tropical species if they facilitate greater investment in physiological adaptations to combat oxidative damage in the face of similar metabolic rates. Quantifying interspecific and latitudinal variation in physiological mechanisms capable of mitigating oxidative damage may reveal how tropical and south temperate species maintain low adult mortality without a major reduction in basal metabolic rate. Mortality rate differences among latitudes may also be due to variation in extrinsic mortality but quantifying latitudinal differences in cause-specific mortality are necessary to test this hypothesis. Overall, our results suggest an urgent need to carefully examine alternative physiological and ecological mechanisms shaping global variation in demographic rates and life histories.

## Supporting information

BMR Survival Literature Data

## ACKNOWLEDGMENTS

We are grateful to Adam Mitchell and Riccardo Ton for comments that improved the manuscript. We would like to thank Alim Biun, Dr. Maklarin Lakim, Rimi Repin and Fred Tuh from Sabah Parks, and C.Y. Chung from the Sabah Biodiversity Centre for assistance in Malaysia. AJB would also like to thank Bonifatreno Jurunin, Enroe bin Soudi, Rayner Ray, Ed Conrad and William Talbot for invaluable assistance in the field. This work was supported by National Geographic Society (9875-16) and the National Science Foundation (DEB-1241041, DEB-1651283 & IOS-1656120) to TEM, an AOU Graduate Student Research Grant and a Wesley M. Dixon Memorial Fellowship to AJB. This work was performed under the auspices of University of Montana IACUC protocol #059-10TMMCWRU. Any use of trade, firm, or product names is for descriptive purposes only and does not imply endorsement by the U.S. Government.

